# A gene program dictionary of human cells

**DOI:** 10.1101/2025.10.29.685322

**Authors:** Yupu Xu, Yifan Wang, Zhenxing Geng, Yue Qin, Shisong Ma

**Affiliations:** MOE Key Laboratory for Cellular Dynamics, School of Life Sciences, Division of Life Sciences and Medicine, University of Science and Technology of China, Innovation Academy for Seed Design, Chinese Academy of Sciences, Hefei, China; School of Data Science, University of Science and Technology of China, Hefei, China

## Abstract

Defining all human cell types and their roles in health and disease is a central goal of biology. Single-cell RNA sequencing has enabled the construction of organ-specific cell atlases, but building a comprehensive organism-wide atlas spanning multiple organs remains challenging due to batch effects, study biases, and inter-organ complexity. Here, we present Gene Program Dictionary (GPD), a framework that leverages robust gene co-expression programs—rather than direct cell integration—to overcome these barriers. Using SpacGPA, a partial correlation-based network method, we analyzed 466 scRNA-seq datasets, generating 1,975 independent networks and 90,701 gene co-expression modules, which were consolidated into 1,534 consensus gene programs representing a wide range of human tissues and cell types. Each program serves as a composite marker, capturing both cell-type-specific and shared biological processes. We demonstrate their utility by mapping endothelial cell subtypes across tissues to reveal their heterogeneity—including tumor-specific programs—annotating colorectal cancer spatial transcriptomes, and linking programs and their corresponding cell types to disease loci, revealing hotspots such as neuronal programs in psychiatric disorders and a proximal tubule program in kidney diseases. GPD provides an organism-wide reference for studying cellular diversity and disease mechanisms.

## Introduction

A central goal of cell biology research is to identify all human cell types and elucidate their roles in health and disease. Single-cell RNA-sequencing (scRNA-seq) has revolutionized this endeavor. By integrating scRNA-seq datasets from diverse laboratories, comprehensive cell atlases have been constructed for individual human organs, such as the brain, lung, eye, kidney, and gut (Elmentaite et al., 2021; Jorstad et al., 2023; Klötzer et al., 2025; Li et al., 2026; Sikkema et al., 2023; Siletti et al., 2023). These atlases are invaluable for understanding organ-specific cellular diversity, identifying marker genes, and mapping lineage relationships. They provide essential references for studying cellular functions and disease mechanisms (Hrovatin et al., 2024).

Despite these advances, constructing a unified, cross-organ cell atlas for the entire human body remains a formidable challenge. This effort is hindered by batch effects across datasets, study-specific biases, and the vast diversity of cell types across organs. While organ-specific atlases mitigate some of these challenges using integration methods such as scANVI, Scanorama, and scVI (Hie et al., 2019; Lopez et al., 2018; Xu et al., 2021), these limitations are magnified when scaling to an organism-wide atlas. As an alternative, cross-tissue atlas focusing on specific cell types—such as myeloid cells, T cells, B cells, and fibroblasts—have been developed (Cheng et al., 2021; Gao et al., 2024; Yang et al., 2024b; Zheng et al., 2021). These atlases have uncovered diseases-associated cell subtypes, underscoring the potential utility of a comprehensive whole-body atlas.

Beyond atlas construction, identifying shared gene programs provides a complementary approach for integrating single-cell datasets. Gene programs are sets of co-regulated genes fulfilling specific biological functions and exhibiting coordinated expression. Non-negative matrix factorization (NMF) is a widely used method to identify gene programs (Kotliar et al., 2019; Lee and Seung, 1999). By applying NMF, a single-cell gene expression matrix can be decomposed into two non-negative matrices that represent gene programs and their usage across cells. This approach has identified gene programs across tumor scRNA-seq samples, which were then integrated into consensus meta-programs to reveal tumor-related pathways with both common and heterogeneous features (Barkley et al., 2022; Gavish et al., 2023; Yang et al., 2024a). A similar strategy was also used to identify reproducible programs underlying T cell subsets (Kotliar et al., 2025). Notably, these studies demonstrated that gene programs, unlike individual cells, are less susceptible to batch effects and are therefore easier to integrate across datasets. Other matrix decomposition-based methods, such as Spectra and MetaTiME, have also been developed for gene programs identification (Kunes et al., 2023; Zhang et al., 2023). However, these approaches have predominantly been applied to tumor datasets, and their use in broader organ-scale atlases remains largely unexplored.

Another approach to uncovering gene programs involves gene co-expression network analysis followed by gene module detection. However, the sparsity of scRNA-seq datasets makes it challenging to infer co-expression relationships. To address this, methods tailored for single-cell gene co-expression network analysis, such as hdWGCNA and SingleCellGGM, have been developed (Morabito et al., 2023; Xu et al., 2024). Among them, SingleCellGGM, recently introduced by our lab, employs partial correlation coefficients (*Pcor*) to evaluate co-expression relationships. *Pcor* measures the correlation between two genes while controlling for the influence of all other genes, making it a robust metric for network analysis. When applied to mouse single-cell datasets, SingleCellGGM successfully identified robust gene co-expression modules, capturing cell type-specific functions, developmental processes, and shared cellular mechanisms, even in the context of dataset sparsity (Xu et al., 2024). Recently, we further optimized the algorithm and developed the SpacGPA framework, featuring GPU-accelerated network construction and gene module identification, enabling efficient processing of large-scale datasets (Xu et al., 2025).

In the present study, we developed a Gene Program Dictionary (GPD) framework that aims to systematically identify gene programs across nearly all human cell types. We reasoned that constructing an organism-level atlas by integrating individual cells from datasets spanning different organs and studies remains impractical due to batch effects, study-specific bias, and inter-organ complexity. To overcome these challenges, we leveraged the robustness of gene programs against batch effects. Using SpacGPA, we identified gene co-expression modules from individual datasets and integrated them to generate consensus gene programs. By processing diverse single-cell datasets from various human organs, we established a GPD encompassing 1,534 robust gene programs. These programs serve as composite markers for cell type identification, including in high-resolution spatial transcriptomic datasets, effectively acting as proxies for individual cell types. Furthermore, the GPD facilitates the discovery of gene programs and cell types associated with human diseases, including “hotspot” programs and cell types implicated in multiple diseases. These findings underscore the GPD’s potential to advance our understanding of cellular mechanisms in health and disease.

## Results

### A GPD framework to integrate human single cell datasets

We developed a GPD framework to identify gene programs across a wide range of human cell types at the whole-organism level (**Figure 1A**). This involved integrating public single-cell transcriptome datasets from a wide range of studies. To mitigate batch effects, we focused on integrating signature gene modules identified from each dataset rather than individual cells.

**Figure 1.**
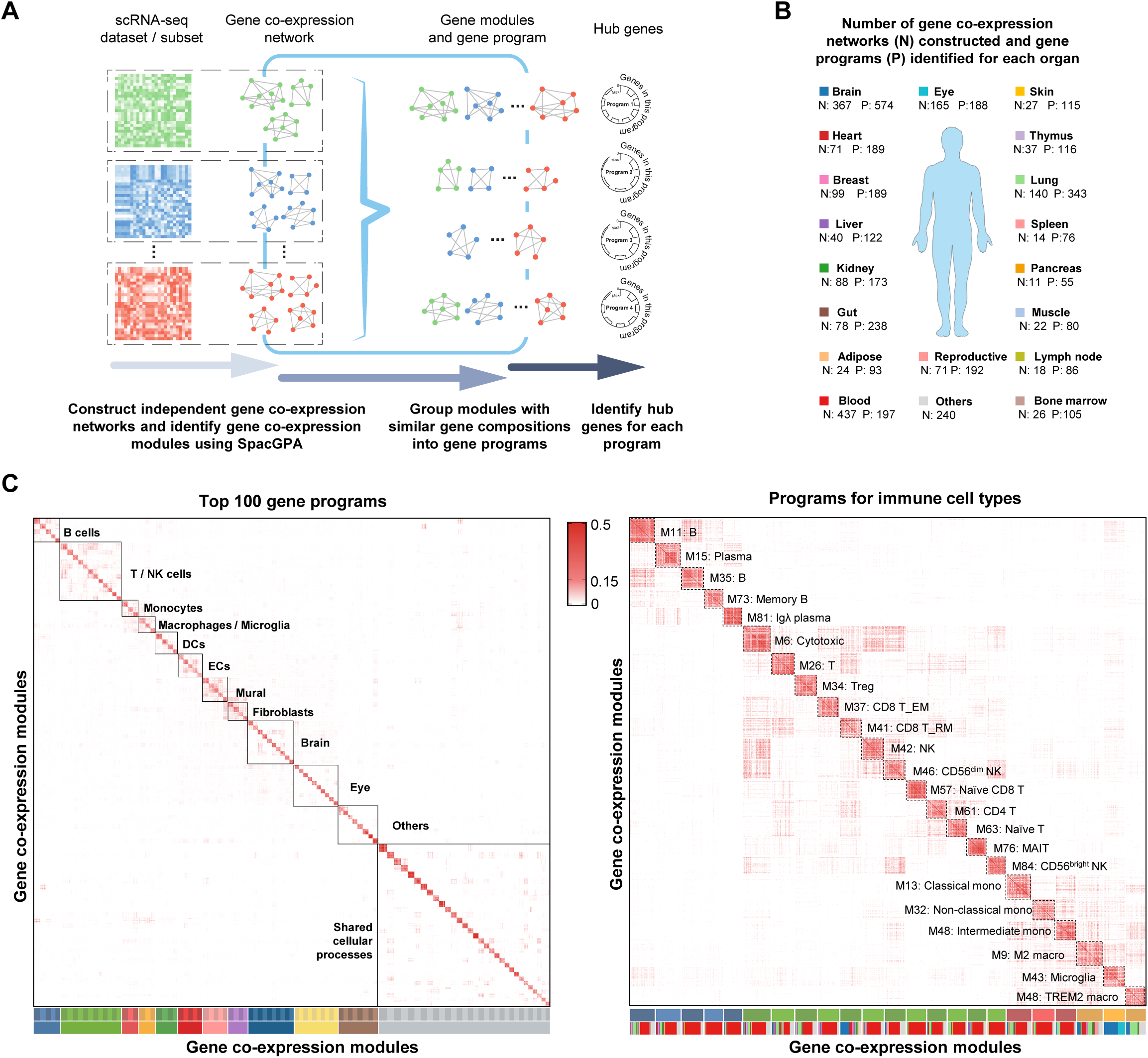
Building a gene program dictionary for human cells. **(A)** The analysis work flow. **(B)** The number of networks constructed and gene programs identified for each organ. **(C)** The top-100 gene programs. Similarities between the constituent modules, quantified by the modified Jaccard index (*Jm*), are shown. Modules were grouped by their corresponding programs, and programs were arranged according to their broad categories. On the left is an overview of the top 100 programs, and on the right is a detailed view of immune-related programs. The tissue origin of the modules are shown at the bottom for the immune-related programs, with colors corresponding to those in (B). Only a subset of the constituent modules is displayed due to space constrains.

We processed almost all human scRNA-seq datasets from the CZ CELLxGENE Discover Census database (Abdulla et al., 2025). The datasets are organized into collections in the database, with each collection, containing one or multiple datasets, considered an independent study. For some datasets, we constructed a gene co-expression network for each dataset using all cells as inputs via the SpacGPA algorithm. For other datasets–often comprising cells integrated from multiple samples–we divided the cells within each dataset non-redundantly into independent subsets based on sample identifiers (e.g., donor IDs), biological characteristics (e.g., age groups), or both, and constructed gene co-expression networks for each subset. The resulting networks were then used to identify gene co-expression modules separately for each network. In total, we analyzed 466 datasets from 196 collections, yielding 1,975 independent gene co-expression networks representing various organs, including 367 for the brain, 437 for blood, and 77 for the gut, leading to the identification of 90,701 gene co-expression modules in total (**Figure 1B; Table S1**).

We next integrated these gene co-expression modules to construct a comprehensive GPD for human cells. Since cell types were typically represented across multiple datasets or subsets, their corresponding gene co-expression modules were identified in multiple networks, often sharing similar gene compositions. To integrate these modules, a modified Jaccard similarity index (*Jm* index; see Methods) was calculated between modules based on their gene compositions. Module pairs with a *Jm* index of 0.15 or higher were connected to form a gene module network. Using the MCL clustering algorithm (Enright et al., 2002), we clustered this network and identified 1,534 module clusters, each comprising at least five modules derived from five different co-expression networks (**Table S2**). These clusters contained 66,960 of the identified modules, while the remaining modules belonged to smaller clusters not considered hereafter. For each cluster, constituent modules were merged to derive a unified gene program, leading to the identification of 1,534 robust gene programs for human cells (**Table S3**). The programs were subsequently annotated for cell-type specificity and functional relevance based on their expression patterns, marker genes, and enriched Gene Ontology (GO) terms (**Table S4 and S5**). As examples, Figure 1C illustrates the top 100 programs, organized by categories, along with the *Jm* indices between their constituent modules, with immune-related programs highlighted in detail.

Since the programs were integrated from datasets across 196 collections, we assessed whether they were influenced by potential batch effects via examining the distribution of constituent modules across collections. Modules from all 1,534 programs were projected into UMAP space and colored by program identity, collection ID, tissue origin, and broad cell type (**Figure 2**). The plots showed that modules from different collections were well mixed within individual programs (**Figure 2A, B**), indicating that clustering was not driven by collection-specific batch effects. Consistent with this observation, 1,351 of the 1,534 programs (88%) contained modules derived from at least two collections, demonstrating high reproducibility across independent studies (**Table S2**). The remaining 183 programs, primarily originated from large eye, brain, breast, and kidney atlas collections, likely reflect collection-specific cell types rather than technical artifacts. For example, the 29 programs restricted to an eye atlas (Li et al., 2026) predominantly exhibited cell-type-specific program expression (defined below) (**Figure S1**), consistent with biologically meaningful specialization.

**Figure 2.**
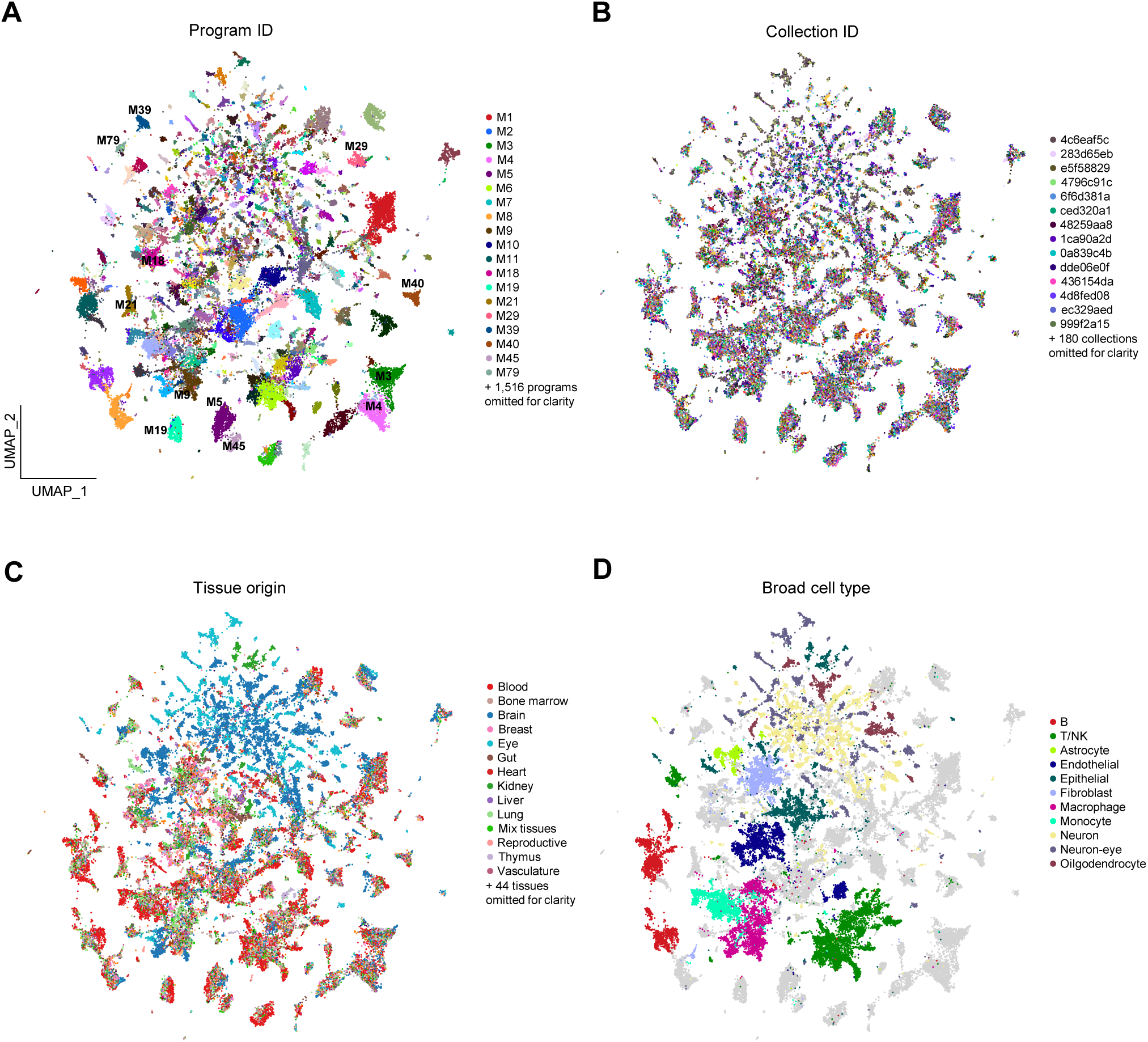
Gene programs are not driven by batch effects. UMAP visualization of constituent modules from all 1,534 gene programs, colored by program ID **(A)**, collection identity **(B)**, tissue origin **(C)**, and broad cell-type categories **(D)**. Each collection was treated as a distinct batch. Each dot represents a constituent module modeled as a virtual cell, in which gene connectivity (degree with the module) was used as the expression value. A kNN graph was constructed in Scanpy using its native Jaccard metric and used for UMAP embedding. Due to space constraints, legends display only a subset of program ID, collections ID, tissues, and broad cell-type categories. For collections, only the first seven characters are shown (full names are 36 characters). In (B), modules from different collections are well mixed within individual programs, indicating that batch effects do not dominate program structure. In (C), both programs shared across diverse tissues and programs with strong tissue specificity are observed. In (D), modules of the same broad cell-type categories cluster in close proximity.

In addition, our framework identified programs representing shared cell types or pathways across multiple tissues as well as programs for tissue-specific cell types; it also brought together programs for cell types belonging to the same broad categories. As shown in Figure 2C, a large portion of the programs contained modules derived from diverse tissues, such as M9 for M2 macrophage, M29 for mast cells, M18 for smooth muscle cells, and M3 and M4 for cell cycle regulation. At the same time, other programs exhibited strong tissue specificity, particularly those for neurons from both brain and eyes. Additionally, programs corresponding to the same broad cell type categories clustered locally in the UMAP plot, including those for B cells, T/NK cells, endothelial cells, epithelial cells, fibroblasts, and neurons (**Figure 2D**). Thus, our framework identified both shared and tissue-specific gene programs across tissues while preserving the overall hierarchical structure of cell types.

Having established the robustness of the framework, we examined individual programs in details. Program M19 comprises 618 gene modules identified across 558 gene co-expression networks, which exhibit substantial overlap in their gene compositions. The repeated identification of these modules underscores their robustness. Three representative modules from program M19 showed specific expression patterns in plasmacytoid dendritic cells (pDC) across diverse datasets (Hao et al., 2021; Perez et al., 2022; van der Wijst et al., 2021) (**Figure 3A**). By integrating these modules, we defined a gene list for program M19 comprising 345 genes. These genes were ranked by their frequency of occurrence across constituent modules, with the most frequently occurring genes designated as hub genes (**Figure 3B**). Notably, these hub genes, considered as the most robust genes within the program due to their repeated occurrence, also serve as marker genes for pDC. Four of the top ten hub genes in M19—*LILRA4*, *CLEC4C*, *IL3RA*, and *LAMP5*—are established pDC markers (Ngo et al., 2024; Villani et al., 2017). Consistent with its constituent modules, the program expression, defined as the frequency-weighted average expression of its top 30 genes, also exhibited specific expression in pDCs, leading to the annotation of M19 as a pDC program. The circo diagram in Figure 3B visualizes this program by displaying its hub genes and their frequencies of occurrence.

**Figure 3.**
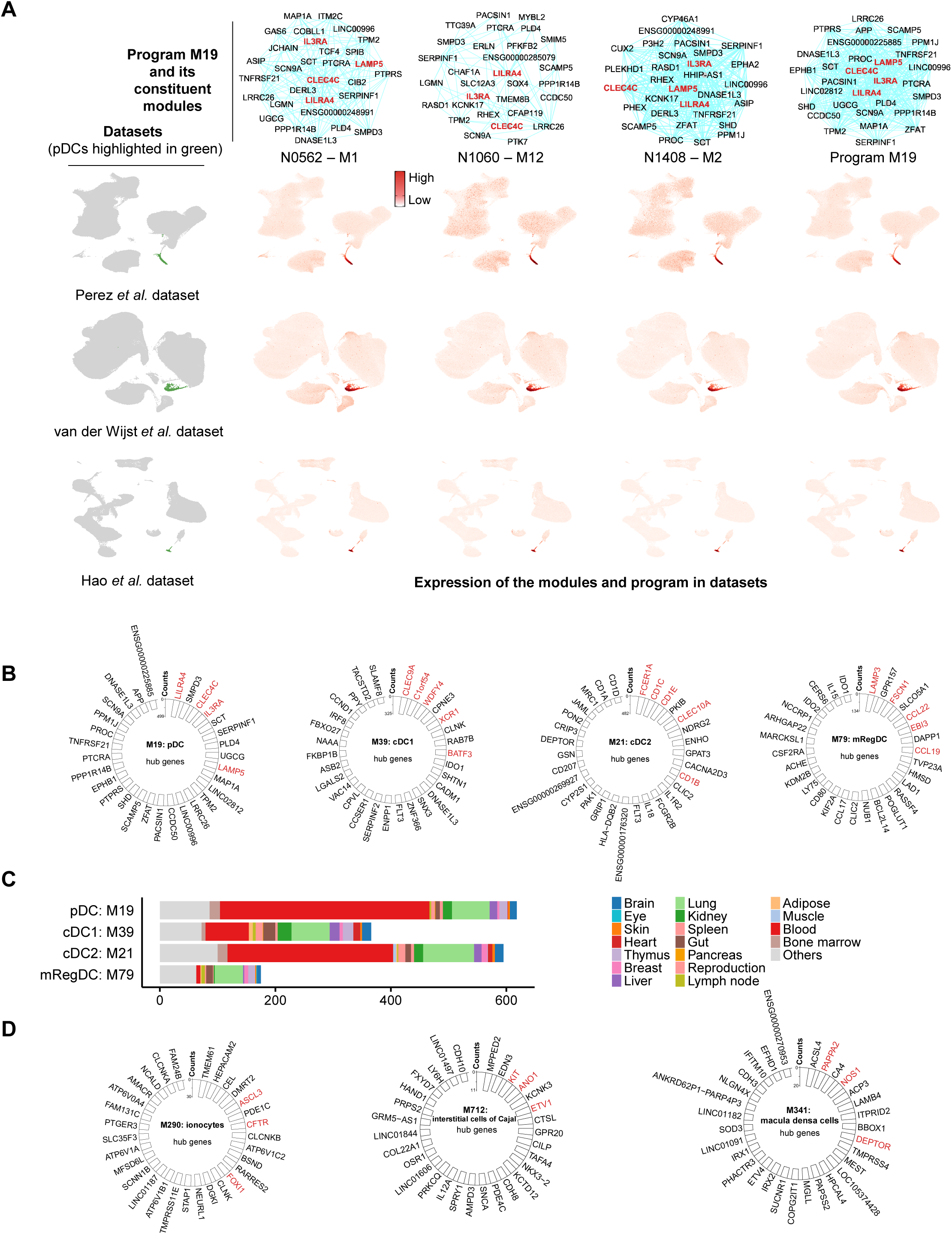
Gene programs for dendritic cells and selected rare cell types. **(A)** A gene program for pDCs. The top panels show subnetworks of three constituent modules and the integrated gene program (M19). Only the top 30 genes from each sub-network are displayed. Genes highlighted in red are marker genes for the corresponding cell type (here, pDCs). The bottom three rows show UMAP of three training scRNA-seq datasets, with pDCs cell highlighted, and the expression of the constituent modules and the integrated program across three independent datasets, demonstrating specific expression in pDCs. The three constituent modules were derived from networks constructed using subsets of the three training scRNA-seq datasets. **(B)** Circo plots displaying the top hub genes of the programs. The axis indicates the number of networks in which the constituent modules were identified, and the occurrence frequency of the hub genes across these networks is also shown. Labels in red indicate marker genes of the corresponding cell types (applies to all subsequent panels). Gene programs for pDC, cDC1, cDC2, and mRegDC are shown. **(C)** Tissue origins of the constituent modules of dendritic cell gene programs. **(D)** Gene programs identified for selected rare cell types.

Similarly, we identified gene programs for three other types of dendritic cells (DCs): type 1 classical DCs (cDC1, M39), type 2 classical DCs (cDC2, M21), and mature DCs enriched in immunoregulatory molecules (mRegDC, M79) (**Figure 3B**). These programs also included known marker genes among their top hub genes: *CLEC9A*, *C1orf54*, *WDFY4*, *XCR1*, and *BATF3* for cDC1 (M39); *FCER1A*, *CD1B*, *CD1C*, *CD1E*, and *CLEC10A* for cDC2 (M21); and *LAMP3*, *FSCN1*, *CCL22*, *EBI3*, and *CCL19* for mRegDC (M87) (Cheng et al., 2021; Del Prete et al., 2023; Villani et al., 2017). Notably, these programs were derived from modules spanning diverse tissues and datasets, demonstrating that module grouping was driven by cell-type specificity rather than batch effects associated with tissue or dataset origin (**Figure 3C**).

Notably, all four DC programs are among the most prevalent programs, ranking within the top 100 by module number. Consistent with this pattern, the top 100 programs are enriched for those associated with common cell types distributed across multiple tissues, including 12 programs for T and NK cells, 5 for B cells, and 6 for monocytes and macrophages (**Figure 1C**). In parallel, a subset of programs captures cell types with strong tissue specificity, including 10 associated with eye-specific cell types and 9 with brain-specific cell types. Beyond cell-type-specific programs, we also identified programs shared across multiple cell types that reflect conserved biological processes, such as M5 (type I interferon response), M45 (type II interferon response), M4 (cell cycle—G1/S), M3 (cell cycle—G2/M), and M40 (DNA damage response) (**Figure 1C**). These results highlight the ability of our GPD framework to robustly identify both cell-type-specific and cross-cell-type gene programs, providing a valuable resource for understanding cellular diversity and function.

While the top 100 programs mainly represent common human cell types and shared processes, our framework also identified programs corresponding to rare cell types across multiple tissues and organs. These include ionocytes (M290) and neuroendocrine cells (M554 and M1501) in the lung, interstitial cells of Cajal (ICC, M712) and fibroblastic reticular cells (M491) in the gut, and α-intercalated cells (IC-A, M149), β-intercalated cells (IC-B, M235), and macula densa cells (MD, M341) in the kidney (**Table S5**). These programs also contain known marker genes among their hub genes, such as *FOXI1*, *ASCL3*, *CFTR* for ionocytes in M290, *KIT*, *ANO*, *ETV1* for ICCs in M712, and *NOS1*, *PAPPA2*, *DEPTOR* for MDs in M341 (**Figure 3D**) (Elmentaite et al., 2021; Lake et al., 2023; Lee et al., 2017; Montoro et al., 2018; Shroff et al., 2021; Sikkema et al., 2023). Thus, our GPD framework identified gene programs and marker genes for both common and rare cell types across diverse tissues and organs.

### Gene programs delineate diversities among endothelial and other major cell types

Human cell types such as B cells, T cells, fibroblasts, monocytes, and macrophages show extensive variability across tissues and disease states. Previous atlas-building efforts, typically focusing on one cell type at a time, captured such diversity and identified disease-related subtypes (Cheng et al., 2021; Gao et al., 2024; Yang et al., 2024b; Zheng et al., 2021), but these approaches often required stringent sample selection and complex processing, which limited coverage and sometimes distorted biological signals. In contrast, our GPD framework integrates nearly all datasets with sufficient cell numbers, covering tissues across all organs and conditions. This allows us to resolve fine-scale subtype differences while maintaining organism-wide representation. Importantly, our approach simultaneously captures gene programs for all major human cell types (**Figure 2D**), providing a unified resource for healthy and disease biology.

As an example, we identified 90 endothelial cell (EC) programs out of 1,534 in total (**Figure 4A, B; Table S6**). Among them, eleven are considered as general EC programs, as they were reproducibly detected across ⩾5 tissues (**Figure 4C; Table S6**). Program M65, assembled from 212 modules across 205 networks spanning gut, lung, kidney, brain, eye, and other tissues, represents a pan-EC program. It is enriched for pan-EC markers (*CD34*, *VWF*, *PECAM1*, *CDH5*) expressed across all ECs regardless of tissue and vascular subtypes (Goncharov et al., 2020). In contrast, M31 defines lymphatic ECs (LECs), carrying classical LEC markers, including the lineage-defining transcription factor gene *PROX1*, junctional protein gene *STAB2*, and surface receptor genes *CCL21*, *FLT4*, *NRP2*, and *LYVE1* (Takeda et al., 2023). At the same time, M64 specifies venous ECs, with marker genes *ACKR1*, *SELP*, and *SELE*, and M71 corresponds to arterial ECs, containing markers *GJA5*, *EFNB2*, *JAG1*/*JAG2* (Schupp et al., 2021). Finally, program M289 is enriched with BMP signaling genes (*SMAD6/7/9*, *ENG*, *ID1/2/3/4*, *SOX18*, *KLF10*, *GATA*), which regulate EC proliferation, angiogenesis, arterio-venous specification, and stress responses, underlying EC plasticity and adaptability. The program thus may represent a signaling-activated, stress-responsive EC subtype commonly seen during vascular remodeling, inflammation, or early angiogenic activation.

**Figure 4.**
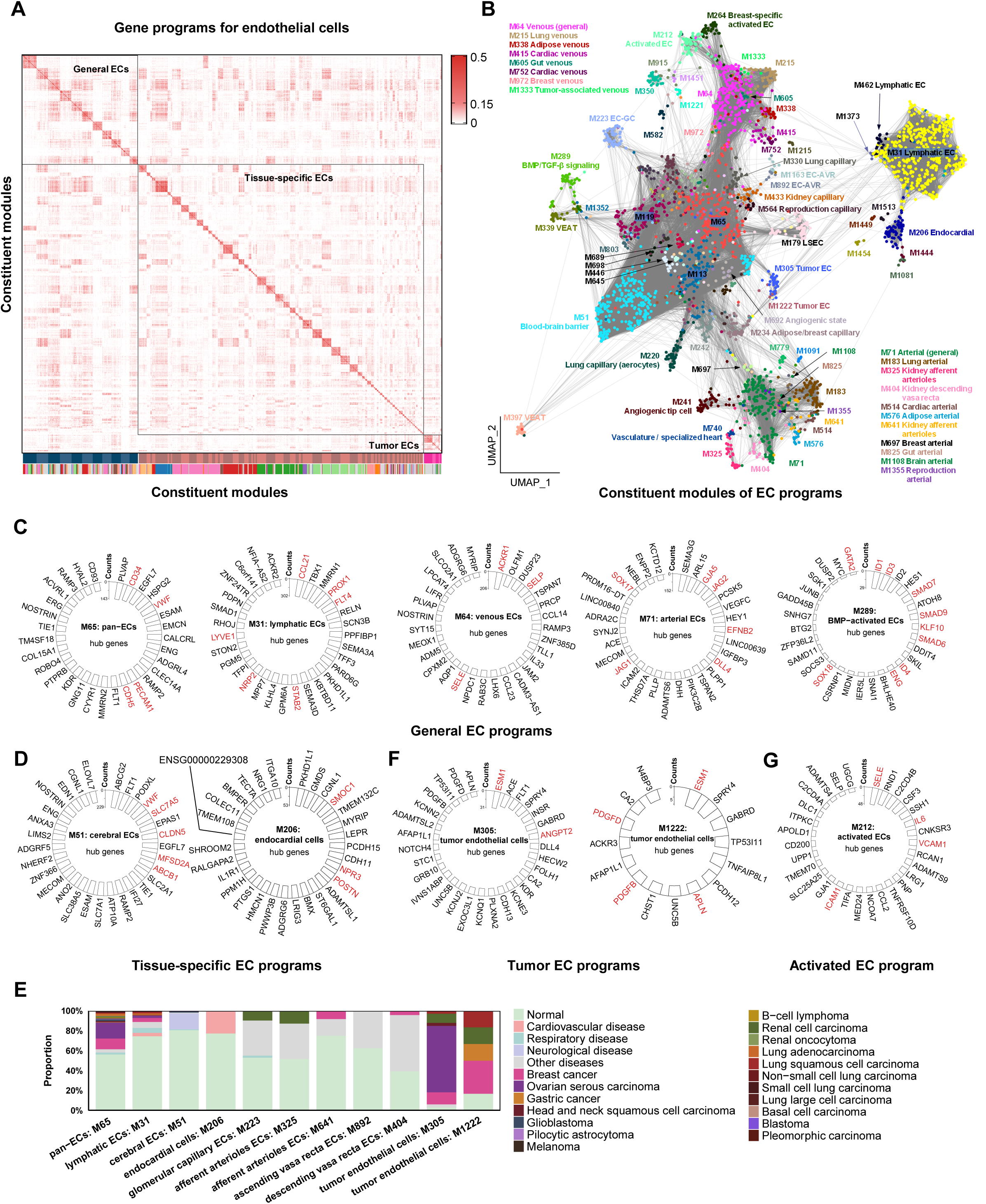
Gene programs identified for endothelial cells across tissues and organs. **(A)** Pairwise similarities among the constituent modules of the 90 endothelial cell gene programs. **(B)** UMAP visualization of the constituent modules from EC programs. Each dot represents a gene module and is colored by program ID. A kNN graph was constructed in Scanpy using a cosine similarity metric between modules and used for UMAP embedding. Modules with *Jm* ⩾ 0.15 were connected by edges to highlight modular structures. **(C)** General EC gene programs shared across diverse tissues. **(D)** Tissue-specific endothelial gene programs. **(E)** Disease origin of selected endothelial gene programs. **(F)** Gene programs for tumor endothelial cells. (**G**) A gene program for activated EC cells.

Besides the general programs, most EC programs exhibit tissue specificity (**Figure 4A, B, D**). For example, M51 is assembled from 265 brain and 5 eye modules and defines cerebral ECs. While containing pan-EC markers (*VWF*, *PECAM1*, *CD34*), it is also enriched for regulators of blood-brain barrier (BBB) functions, such as *CLDN5*, *MFSD2A*, *ABCB1*, *ABCG2*, and *SLC7A5*, which either maintain BBB integrity or regulate transport across BBB (Ronaldson and Davis, 2024; Wälchli et al., 2024). Another program, M206, marks endocardial cells (EndoCs), specialized ECs lining heart chambers and valves that are critical for heart development, signaling, barrier function, and repair. This program includes canonical EndoC markers like *SMOC1*, *NPR3*, and *POSTN* (Reichart et al., 2022). It also contains genes regulating adhesion, ECM remodeling, immune modulation, stress responses, and secreted/paracrine factors, serving as ideal candidates to study heart development and diseases. We also identified EC programs with specific expression in breast, kidney, lung, liver, skin, and reproduction organs (**Table S6**). For example, in kidneys, ECs of glomerular capillary (EC-GC, M223), afferent arterioles (EC-AEA, M325, M641), ascending vasa recta (EC-AVR, M892), and descending vasa recta (EC-DVR, M404) were identified. Across tissues such as lung, heart, breast, adipose, and gut, tissue-specific programs for capillary, venous, and arterial endothelial cells were also identified. In UMAP space, these programs localized near each other according to sub-cell-type identities rather than tissue specificity, indicating that the GEPs capture both shared and subtle differences among subtypes (**Figure 4B**). These programs thus dissect the vast diversity of ECs across healthy human tissues.

Our framework also revealed EC programs strongly enriched in disease contexts, particularly tumors. While many EC programs arose from both healthy and diseased samples—for example, M31 (LEC) and M65 (general EC) contained 24–36% tumor-derived modules—two programs stood out as highly tumor-biased: M305 and M1222, with 97% and 83% of modules derived from tumors, respectively (**Figure 4E**). M305 appeared in ovarian, renal, breast, and other cancers, while M1222 was detected in breast, gastric, lung, and kidney cancers. This disproportionate enrichment indicates that these programs define tumor endothelial cells (TECs). Consistently, their hub genes highlight canonical tumor-associated angiogenic mechanisms (**Figure 4F**). In M305, the top hub gene is *ESM1*, encoding a secreted protein that potentiates VEGF-driven angiogenesis by binding VEGF-A and VEGFR2, thereby enhancing sprouting angiogenesis (Rocha et al., 2014). *ESM1* is broadly upregulated in TECs across solid tumors, with elevated levels linked to metastasis and poor prognosis (Sarrazin et al., 2006). Another hub gene, *ANGPT2*, destabilizes vessels and acts as a pro-angiogenic switch, promoting vascular sprouting, leakiness, and recruitment of tumor-promoting immune cells (Akwii et al., 2019). ANGPT2 is also a key factor limiting VEGF-targeted therapies, as its upregulation drives resistance (Rigamonti et al., 2014). M1222, while also centered on *ESM1*, further contains angiogenic drivers such as *APLN*, *PDGFB*, *PDGFD*, underscoring the multifaceted pro-angiogenic programs active in tumors (Andrae et al., 2008; Uribesalgo et al., 2019). Together, these two tumor-specific programs highlight both well-established and novel TEC regulators, providing ample candidates for biomarker discovery and therapeutic targeting.

Beyond normal EC and TEC-associated programs, our framework also identified M212, which corresponds to ECs undergoing activation and inflammatory stress responses (**Figure 4G**). This program contains hallmark genes of EC activation and cytokine signaling, including *SELE*, *VCAM1*, *ICAM1*, and *IL6*. Collectively, these results demonstrate that our framework captures EC programs spanning baseline, activated, and pathological states, offering a comprehensive view of EC function across diverse physiological and disease contexts.

In addition to ECs, our GPD framework also identified programs for other major cell types, including 87 for fibroblasts, 33 for B cells, 22 for astrocytes, 174 for epithelial cells, 111 for monocytes and macrophages, 98 for T/NK/NKT cells, 24 for oligodendrocytes, and 285 for neurons, spanning both healthy and diseased states (**Figure 2D, Table S6**). Together, these programs offer a broad foundation for dissecting cellular diversity across the human organism.

### Gene programs facilitate annotation of novel spatial transcriptome data

The 1,735 identified gene programs span a wide range of human organs. A program was considered present in a given organ if its constituent modules were derived from two or more co-expression networks constructed from datasets of that organ. In total, 197 genes programs were identified in blood, 105 in bone marrow, 574 in brain, 189 in breast, 238 in gut, and 173 in kidney, among others (**Figure 1B; Table S7**). Within each organ, these gene programs function as composite signature markers that distinguish specific cell types and states, thereby facilitating the annotation of new single-cell and spatial transcriptome datasets.

As an example, we used the 238 gut programs as a reference to annotate an independent spatial transcriptome dataset generated via the 10x Visium High Definition (HD) platform, which profiled genome-wide spatial gene expression in a colorectal cancer (CRC) sample (P2; **Figure 5A**) at a 2 μm resolution (Oliveira et al., 2025). For this analysis, we utilized the 16 μm resolution, comprising 126,768 gapless 16 μm-binned square spots across a 6.5 × 6.5 mm area after filtering out low-expression spots. Using SpacGPA, we constructed a gene co-expression network for the dataset and identified 65 gene modules, each considered as a program (N1-N65; **Tables S8**). To annotate these programs, we compare them to the 238 reference programs and assigned annotations based on the most similar matches. For example, program N28 is most similar to reference program M123 (*Jm* = 0.347 and *P* = 2.12e-68) and was annotated as a neutrophils program (**Table S9**). The annotation is further supported by the presence of neutrophil markers (*S100A8*, *S100A9*, *FCGR3B*) among N28’s hub genes. Using a similar strategy, 39 of the 65 novel programs were annotated (*Jm* ⩾ 0.1 and *P* ⩾ 1e-8; **Table S9**). For another 6 programs without qualified matches, the best hit reference programs also provide clues and aide in their annotations, which were further confirmed by their expression in independent test datasets (Elmentaite et al., 2021; Nowicki-Osuch et al., 2023) or marker genes they contained, such as program N3 for BEST4+ epithelial, N20 for MMP9+ macrophage, and N54 for CCL11+ fibroblast. Among the rest un-annotated programs, a large portion are associated with CRC tumor cells, as demonstrated below.

**Figure 5.**
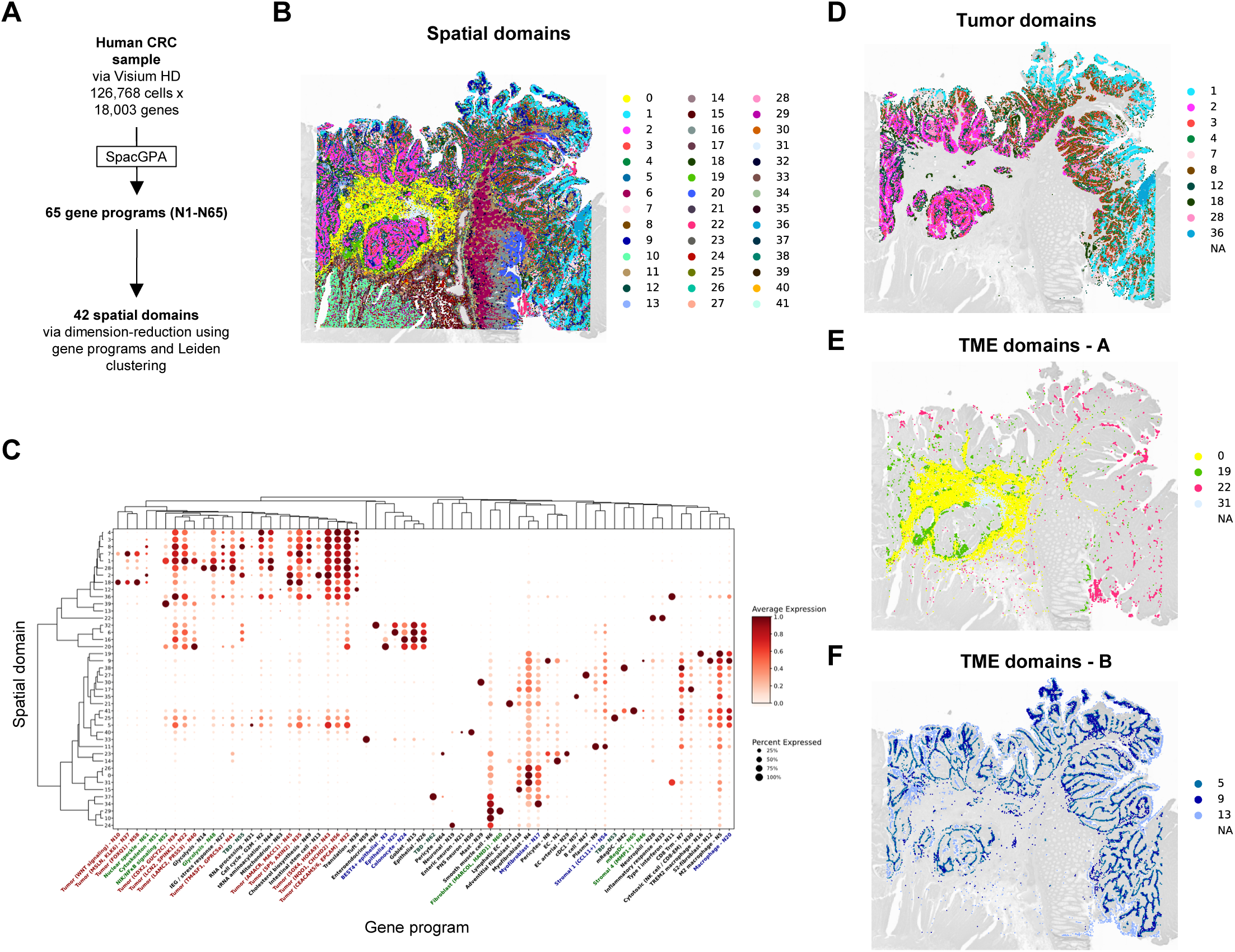
Annotation of a novel CRC Visium HD sample using the reference gene programs. **(A)** Identification of 65 novel gene programs (N1-N65) from the CRC sample. **(B)** Spatial domain identification via Leiden clustering of spots based on gene program x spots dimension reduction. **(C)** Dot plot heatmap showing domain-specific expression of the novel gene programs. Programs are colored according to whether they were annotated using the reference dictionary. Refer to Table S9 for details. **(D)** Tumor-specific spatial domains. **(E, F)** The tumor micro-environment.

These annotated programs can be used for directly annotating spatial spots, highlighting domains for neutrophils, mast cells, goblet cells, colonocyte, BEST4+ epithelial cells, mRegDCs, and others (**Figure S2**). Alternatively, we annotated spots using clustering based on program-driven dimensionality reduction. Program expression per spot was calculated, and the normalized spot × program matrix was used to construct a KNN network, followed by Leiden clustering to group spots into 42 spatial domains (**Figure 5B**). A heatmap was constructed to group spatial domains and gene programs with similar expression patterns together, using functions provided by SpacGPA, which illustrates the expression level of each program with each spatial domain (**Figure 5C**).

Ten spot clusters (#4, #3, #8, #7, #1, #28, #2, #18, #12, #36) were identified as tumor clusters based on copy number variation and marker gene expression (**Figure 5D**). These clusters grouped together in the heatmap and expressed hallmark tumor programs, such as cell cycle (N2) and tRNA aminoacylation programs (N44). They also shared six programs —N22, N32, N34, N35, N43, and N56 —in common (**Figure 5C**). Among them, N32 showed limited similarity to a reference epithelial program (*Jm* = 0.097, M78) and contained CRC marker genes *CEACAM5*, *CEACAM6*, and *EPCAM*, thus serving as a composite marker for CRC (Goossens-Beumer et al., 2014; Oliveira et al., 2025). Programs N22, N34, N35, N43, and N56 were novel, not matched to the reference, and each contained CRC markers *LCN2*, *GUCY2C*, *AXIN2*, *SOX4*, and *NQO1*, respectively (Andersen et al., 2009; Chaudhary et al., 2021; Ji et al., 2014; Lisby et al., 2021; Yan et al., 2001). Additional programs highlighted intra-tumor heterogeneity: Cluster #12, expressing program N38 involved in translation, marked the tumor core facing the luminal, while Cluster #18 represented the invasive front between tumor and normal adjacent tissue (NAT), specifically expressing program N10, associated with WNT signaling (*NOTUM*, *WNT6*, *NKD2*) and critical for CRC progression (Zhao et al., 2022). Moreover, Clusters #2, #8, #28, #1, and #36 corresponed to distinct tumor regions (from left to right in **Figure 5D**), possibly reflecting different progression states. These clusters exhibited relatively higher expression of programs N13 (intestinal stem cell), N49 (cholesterol biosynthesis), N14 (glycolysis), N22, and N11 (type I interferon response), respectively (**Figure 5C**). Thus, several programs not annotated by the reference set were nonetheless associated with CRC, including N10, N22, N32, N34, N35, N43, N56, and additional programs N40 and N41. This may reflect the limited number of CRC samples used to define the reference gene programs.

The clusters and programs also delineated tumor microenvironments (TME; **Figure 5E**). Cluster #22, expressing a neutrophil signature program (N28) and an inflammatory response program (N33), appeared in the luminal TME and was considered tumor-associated neutrophil (TAN), consistent with high expression of inflammatory hub genes *IL1B* and *CXCL8* in N33. Surrounding tumor Cluster #2 were fibroblasts and macrophages clusters: Cluster #0 expressed myofibroblast program N4 (*COL1A1*, *ELN*, *VCAN*, *MMP2*, *MMP11*) and was considered myofibroblastic cancer-associated fibroblasts (myCAFs); Cluster #31 expressed N4 along with type I interferon response program N11 and was classified as ISG+ myCAFs; Cluster #19 expressed macrophage programs N5 (M2 macrophage) and N16 (*CHIT1*, *SPP1*, *APOE*, *GPNMB*), identified as CHIT+ SPP1+ tumor-associated macrophages (TAMs). Interestingly, these myCAF and TAM domains (#2, #19, #31) did not overlap with TAN domains (#22), warranting further investigation into potential mutual exclusion.

Three additional spatial domains were identified on opposite sides of the tumor: Cluster #13 in the luminal TME and Clusters #5 and #9 at the tumor edge (**Figure 5F**). Cluster #13 did not show clear gene expression patterns except for high expression of a mitochondrion-related program (N63). Cluster #5, expressing program N31 with genes involved in RNA processing and microtubule-based vesicle transport, aligned closely with the tumor edges and may mediate interaction between tumor and TME. In contrast, Cluster #9, located outside of Cluster #5, highly expressed a fibroblast program (N12) and a macrophage program (N20; markers *MMP12*, *MMP10*, and *PLA2G*), indicating that Cluster #9 is a spatial domain composed of fibroblasts and macrophages,. N12 shared similarity with the S2-fibroblast reference program (M492) and included ligands in WNT/EMT/PDGF/growth factor pathways (*WNT4/5A/5B*, *BMP5/7*, *TGFB2*, *CCN4*, *DLL1*, *PDGFD*, *NRG1*, *ANGPTL4*, *APELA*, *CSF1*) as well as ECM/matricellular factors affecting EMT (*EDIL3*, *EMILIN1*, *ADAM19*, *ADAMTS8/14*, *FBN2*, *LAMB1*, *LAMC1*). Thus, S2- fibroblasts in Cluster #14 may provide ligands that influence EMT and CRC tumor progression. Indeed, previous work reporting the S2-fibroblast indicated that they resemble CAFs, are present even in nondysplastic stages of tumor development, and may establish a protumorigenic environment that promotes the emergence of future cancer cells (Nowicki-Osuch et al., 2023).

Overall, gene program-based analysis revealed detailed heterogeneity within CRC tumors and their TMEs, providing much finer resolution than standard pipelines. Reference gene programs facilitated efficient annotation of novel programs and placed them in the broader single-cell context, supporting easier interpretation.

### Gene programs and cell types implicated in human diseases

We next explored the utility of the gene programs in studying human genetic diseases. While numerous disease-associated genetic loci have been identified, the etiological mechanisms underlying most diseases remain unclear. A promising approach involves investigating the convergence of disease loci onto specific biological pathways. However, prior studies have often been constrained by the limited scope of available pathway collections. The extensive coverage of our gene programs offers a valuable opportunity to overcome this limitation. To test this, we obtained significant disease-associated gene loci (*P* ⩽ 5e-8) from the NHGRI-EBI GWAS Catalog for 67 human diseases/traits and analyzed their enrichment within gene programs and corresponding cell types (**Figure 6A, Table S10**) (Sollis et al., 2023).

**Figure 6.**
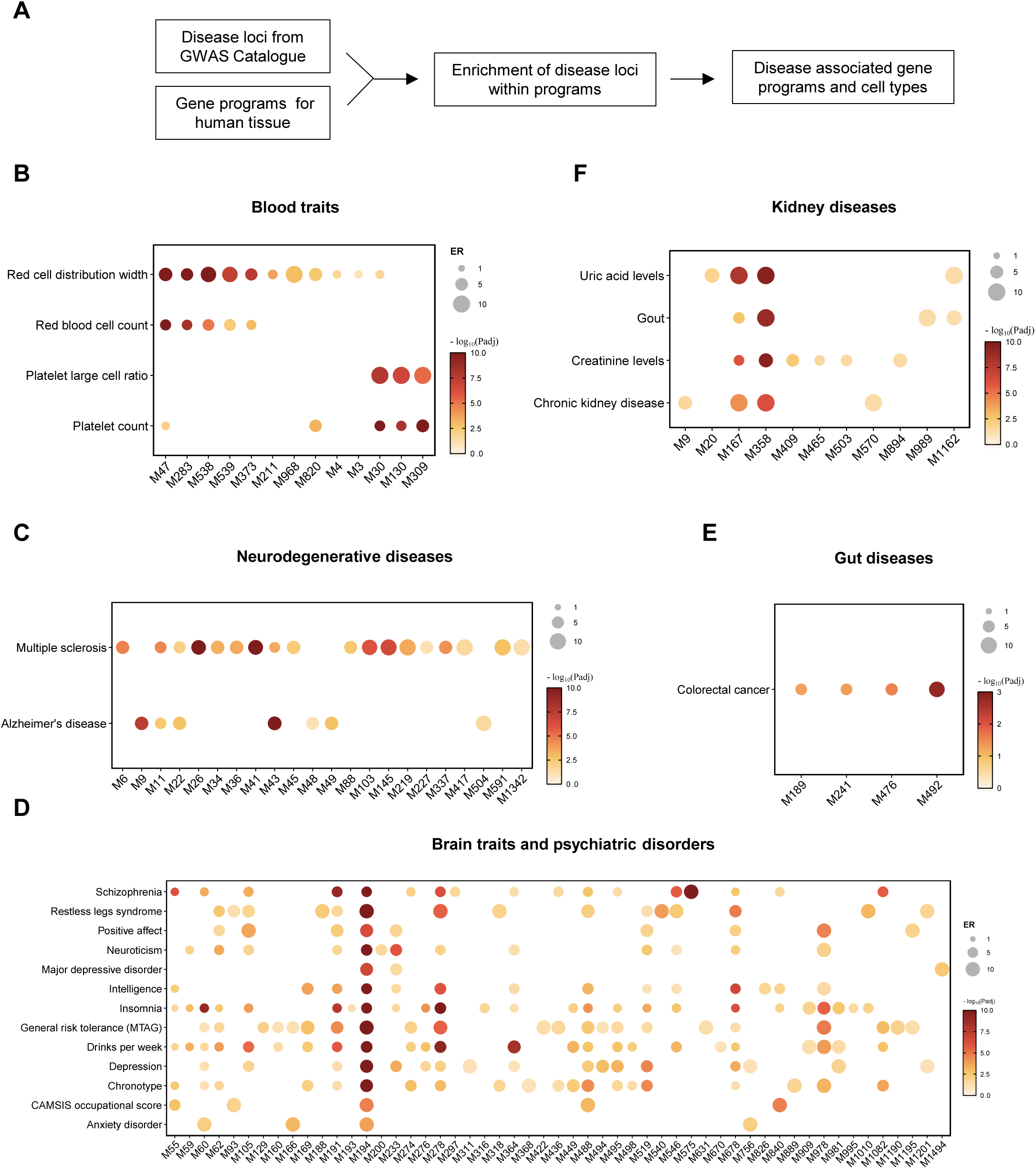
Enrichment of disease gene loci within gene programs. **(A)** The analysis workflow. **(B)** Enrichment of blood-related traits in blood and bone-marrow programs. The color-bar for *P* values is capped at 10. **(C)** Enrichment of neurodegeneration-associated genes within brain programs. **(D)** Enrichment of brain disease genes within brain programs. **(E)** Enrichment of gut-related disease genes within gut programs. **(F)** Enrichment of kidney disease genes within kidney programs.

As a proof of principle, we identified significant enrichment of genes associated with blood traits among the 220 gene programs derived from blood and/or bone marrow (**Figure 6B**). For example, genes linked to red blood cell (RBC) count were enriched within M47, a program specific to erythrocytes. Among the 1,031 genes associated with RBC count, 69 were found in M47, including *ALAS2*, *HBB*, *SLC25A37* and *ANK1*, representing a 3.7-fold enrichment over the background (*Enrichment Ratio [ER]* = 3.7, *P* = 3.65e-19, BH-adjusted). Additionally, three other erythrocyte programs (M283, M538, M539) and another program specific to erythroid progenitor cells, M373, also showed significant enrichment. For red cell distribution width (RDW)—an indicator of anemia—eight erythrocyte-lineage programs (M47, M283, M538, M373, M539, M211, M968, M820) and two additional programs related to DNA replication (M4) and cell cycle regulation (M3) were identified. These findings are consistent with prior studies linking cell cycle dysfunction regulation to conditions such as Fanconi anemia (D’Andrea and Grompe, 2003). Similarly, programs specific to megakaryocytes (M30, M130, M309) were enriched for genes associated with platelet count and platelet large cell ratio. These expected enrichments validate the utility of our approach.

We next assessed enrichment of genes associated with neurodegenerative diseases within the 574 gene programs derived from brain tissues (**Figure 6C**). Consistent with prior studies, genes linked to Alzheimer’s disease (AD) were highly enriched in M43, a microglia-specific program (*ER* = 6.9, *P* = 1.24e-12). This program includes key AD genes such as *ABI3*, *MS4A4A*, *MS4A6A*, *PLCG2*, *TREM2* and *SPI1*. Enrichment of AD-associated genes was also observed in a related program for M2 macrophages (M9, *ER* = 6.4, *P* = 2.46e-08), potentially representing border-associated macrophages. Similarly, genes associated with multiple sclerosis (MS), a neurodegenerative disease characterized by axonal loss, were enriched in the microglia-specific program (M43, *ER* = 3.9, *P* = 1.70e-4), a T cell-specific program (M41, *ER* = 8.1, *P* = 3.65e-14), and an inflammatory response program (M36, *ER* = 6.7, *P* = 1.79e-4). These findings effectively link gene programs and cell types to neurodegenerative disease mechanisms, validating the approach and revealing additional targets for future research.

We further examined enrichment of disease-associated genes for psychiatric disorders and brain-related traits (**Figure 6D**). Disorders analyzed included anxiety disorder, depression, insomnia, major depressive disorder (MDD), neuroticism, schizophrenia, and restless legs syndrome (RLS). Brain-related traits included chronotype, drinks per week, CAMSIS occupational score, intelligence, general risk tolerance (MTAG) and positive affect. Among the 574 gene brain-derived programs, three programs were prominently associated with these disorders and trains. M194 emerged as a pan-neuronal program with widespread expression across all types of neurons and was enriched for genes involved in neuron projection and synaptic transmission (*P* = 8.45e-36 and *P* = 8.34e-34, respectively) (**Figure 6D**). This program showed enrichment for genes associated with all analyzed psychiatric disorders and traits (*ER* = 4.9–10.3, *P* = 1.13e-4–1.55e-28). Importantly, the enriched gene sets varied across disorders and traits, ruling out potential artifacts. M194 thus represents a “hotspot” program for psychiatric disorders and highlights the importance of proper neuron projection and synaptic transmission. A related pan-neuronal program, M191, exhibited similar involvement. Another program, M278, with specific expression in cerebral granular cell, belonging to excitatory neurons, also associated with a number of disorders, highlighting the role of cerebellum in in psychiatric and cognitive phenotypes.

Conversely, programs with more specific expression patterns were linked to fewer disorders. For example, M316, specific to SST interneurons, was associated with insomnia; M169, specific to CGE interneuron, was linked to chronotype, intelligence, drinks per week, and general risk tolerance; and M188, specific to L6b cortical neurons, was associated RLS. Similarly, non-neuronal programs contributed to specific disorders: M55, specific to oligodendrocyte, was linked to schizophrenia, intelligence, drinks per week, chronotype, and CAMSIS occupational score, while M60, specific to oligodendrocyte precursor cells, was associated with insomnia, schizophrenia, general risk tolerance, drinks per week, depression, chronotype, and anxiety disorder. These findings indicate that broadly expressed neuronal programs are linked to multiple brain disorders and traits, whereas programs specific to particular neuron types have specialized roles in certain psychiatric and cognitive phenotypes.

We also analyzed disease gene enrichment for other tissues (**Figure 6E, F**). Among the 238 gut-related gene programs, M492, specific to S2-fibroblasts, was enriched for genes linked to colorectal cancer (CRC) (*ER* = 9.1, *P* = 1.63E-03), with disease genes like *BMP2*, *BMP4*, *BMP5*, and *SOX6*. Spatial transcriptomic analysis hitherto (Figure 4G) has identified S2-fibroblasts with the CRC tumor microenvironment; the enrichment of CRC genes further reinforces their possible role in CRC pathogenesis. Similarly, among the 173 kidney-derived gene programs, M358, specific to proximal tubule epithelial cells (PT), was significantly enriched for genes linked to chronic kidney disease (CKD), gout, and creatinine and uric acid levels. CKD, gout, and uric acid loci showed 27.4-, 10.8-, and 22.2-fold enrichment, respectively (*P* = 4.40e-7, 1.11e-9, and 2.24e-10). These findings align with the established role of PT in uric acid reabsorption and secretion, implicating PT as a vulnerable and “hotspot” disease cell type in CKD pathogenesis.

Similarly, disease gene enrichments were identified for heart, eye, and immune system-related traits and diseases (**Table S10**). Collectively, these results underscore the utility of gene programs in linking disease-associated gene loci to specific cell types and pathways, providing valuable insights into the cellular and molecular mechanisms underlying human diseases.

## Discussion

In this study, we developed a GPD framework to identify gene programs for nearly all human cell types. The resulting 1,534 programs serve as composite markers for both common and rare cell types across all major human tissues. These programs demonstrate robustness, as each comprises at least five constituent modules consistently identified across multiple networks. The recurrent nature of these modules enables the ranking of genes by their frequency of occurrence, facilitating the identification of hub genes within each program. In many cases, these hub genes also serve as marker genes for the corresponding cell types. Importantly, unlike traditional approaches that require prior identification of cell types to define marker genes, our framework identifies gene programs and hub genes independently of cell type annotation. This reverse approach first uncovers cell-type-specific gene programs, which are then assigned to corresponding cell types. This feature is particularly advantageous in scenarios where cell type annotation is ambiguous, challenging, or inconsistent between different studies.

The comprehensive scope of our GPD makes it broadly applicable to a variety of downstream analyses. For example, we successfully applied the gene programs to annotate a novel Visium HD spatial transcriptome dataset. To ensure accurate annotation, our pipeline verifies whether a known gene program can also be identified within the novel sample. Detection of a program in the sample confirms the presence of the corresponding cell type. Additionally, the framework enables the discovery of novel gene programs associated with previously uncharacterized cell types. Gene modules for a novel cell type, when identified across multiple networks more than five times, are clustered into a new module cluster, enabling the confident identification of the corresponding program. Using this approach, we identified 56 known and 16 novel programs in the spatial dataset, facilitating detailed annotation of spatial domains with the sample. Given the broad tissues coverage of our GPD, it is readily extendable to single-cell or spatial datasets from other tissues.

Our GPD framework also offers a powerful tool for studying human diseases. By analyzing the convergence of disease-associated genetic loci onto gene programs, we can pinpoint programs and their corresponding cell types associated with various diseases. The extensive coverage of our gene programs enables such analyses to be conducted across multiple tissues and organs. Notably, our analysis identified “hotspot” gene programs associated with multiple diseases. For instance, the pan-neuronal program M194 was linked to all analyzed psychiatric disorders and brain traits, highlighting its significance in human disease. Rather than being specific to a single neuronal subtype, M194 appears to regulate neuronal projection and synaptic transmission across all neuron types, offering novel insights into the mechanisms underlying psychiatric disorders.

Another example of a “hotspot” program is M358, specific to proximal tubule epithelial cells in the kidney. This program was associated with several interrelated kidney traits, including uric acid levels, creatinine levels, gout, and CKD. These traits are biologically interconnected, as elevated uric acid levels are a primary factor in gout, while creatinine levels serve as a key indicator of CKD (Kuo et al., 2015; Pottel et al., 2016). The association of M358 with these related traits underscores its relevance to kidney disease pathogenesis and further highlights the reliability of our findings. These examples demonstrate the utility of the GPD framework in uncovering disease mechanisms and identifying gene programs of critical importance.

In conclusion, we developed a GPD framework to integrate single-cell transcriptomic datasets from diverse studies and tissues. This framework has successfully constructed a gene program dictionary of human cells comprising 1,534 gene programs. These programs provide an invaluable resource for advancing our understanding of cellular functions in both health and disease.

## Materials and methods

### The GPD framework

To construct a gene program dictionary (GPD) for human cells, we developed a GPD framework. Human single-cell RNA-seq (scRNA-seq) datasets were collected from the CZ CELLxGENE Discover Census database (LTS version 2025-01-30). For part of the datasets, all cells from each dataset were used as an input for single-cell gene co-expression network analysis. For others, datasets were divided into independent subsets according to sample characteristics, with the breakdown keys listed in Table S1. To avoid duplication, only cells labeled as primary data were retained for network analysis. Datasets or independent subsets containing ⩾7,000 cells were treated as independent samples for network construction.

Gene co-expression networks were constructed for each input sample. Raw counts were normalized by total counts (target_sum = 1e4) and log1p-transformed. Genes expressed in fewer than 10 cells were filtered out. The resulting matrix was analyzed using the SpacGPA package (default parameters) to generate gene co-expression networks. A permutation test based on random expression shuffling was performed to estimate the false discovery rate (FDR). If, at *pcor* = 0.02, the FDR was ⩽0.05, gene pairs with *pcor* ⩾0.02 and co-expression in ⩾10 cells were retained for network construction. Otherwise, *pcor* was adjusted to the minimal threshold yielding FDR ⩽ 0.05, and gene pairs meeting these criteria were kept. Networks were clustered using the MCL-Hub algorithm (implemented in SpacGPA) with the parameter inflation = 2, yielding gene co-expression modules. Modules containing ⩾15 genes were retained for further analysis.

In total, 90,701 gene co-expression modules were obtained from 1,975 independent networks. A modified Jaccard similarity index (*Jm*) was calculated to quantify similarity between modules. Because module sizes varied depending on sample cell numbers (larger samples tended to yield larger modules), we standardized module sizes before comparison. For any two modules with *m* and *n* genes, both were trimmed to contain *min(m, n)* genes ranked by intra-module degree, and the top *min(m, n)* genes were used to compute *Jm*. Module pairs with *Jm* ⩾0.15 were used to construct a module network, which was then clustered using the MCL algorithm to obtain meta-modules of gene modules. Meta-modules containing at least five constituent modules derived from at least five independent networks were retained for further analysis.

Meta-modules were integrated to obtain consensus gene programs. For each meta-module, we counted the frequency of gene appearances across constituent modules and ranked genes accordingly. Genes with higher occurrence were considered more stable and treated as hub genes. A permutation test—randomly assigning background genes from each sample to modules—was used to calculate FDRs for each frequency count. Genes with FDR ⩽0.05 and appearing at least three times were retained for the program. Program sizes were capped at the size of their largest constituent module. Consensus sub-networks were constructed by aggregating all edges from constituent modules, with edge weights defined by their occurrence frequency. Programs were annotated by examining their expression in corresponding datasets, performing GO enrichment analysis, and identifying known marker genes.

### UMAP visualization of gene programs

UMAP was used to visualize similarity relationships among constituent modules of gene programs. For each module, a “virtual cell” was constructed in which genes within the module were assigned expression values equal to their connectivity (i.e., the number of edges to other genes within the module), while all other genes were set to zero. An AnnData object was then generated using virtual cells from all modules across the 1,534 identified programs. A kNN graph was constructed from this virtual expression matrix using the Jaccard similarity metric implemented in Scanpy, and subsequently used for UMAP embedding and visualization.

For modules derived from cell types within the same broad cell-type categories, subset AnnData objects were created and visualized separately using UMAP, with similarity measured by the cosine metric. Within these subset visualizations, module pairs with *Jm* ⩾ 0.15 were connected by background edges to further illustrate their relationships.

### Annotating gene programs from novel datasets

To annotate gene programs in a novel dataset, we first constructed a single-cell gene co-expression network and identified gene modules using SpacGPA. These modules were treated as novel gene programs and compared to reference programs using three measures: the Jm similarity index, the P-value of overlap, and the number of shared genes, all based on trimmed modules as outlined above. A composite ranking score was computed by multiplying the Jm rank, the square root of the P-value rank, and the square root of the shared gene number rank. Programs with the lowest composite ranking score were considered the most similar. Significant matches were defined as those with *Jm* ⩾0.1 and *P* ⩽1e−8. As an example, we analyzed a Visium HD spatial transcriptomics dataset from a colorectal cancer (CRC) sample (10x Genomics) and annotated its gene programs using the reference dictionary.

### Disease gene program analysis

To identify enrichment of disease-associated genes within gene programs, we obtained disease loci from the GWAS Catalog (as of 2025-09-15). Only loci with *P* ⩽5e−8 and located within gene bodies were included. Enrichment analysis was performed using a hypergeometric test for diseases and traits related to the blood, brain, gut, kidney, and other tissues.

### Data and code availability

All datasets used in this study are publicly available from CZ CELLxGENE Discover Census, 10x Genomics, and the NHGRI-EBI GWAS Catalog, as indicated in the text. The code for the Gene Program Dictionary (GPD) framework will be made available on GitHub (https://github.com/MaShisongLab/GeneProgramDictionary).

## Acknowledgements

We thank the USTC Supercomputing Center and USTC School of Life Sciences Bioinformatics Center for providing the computing resources.

## Author Contributions

**Yupu Xu:** Methodology, Investigation, Formal analysis, Writing – Original draft. **Yifan Wang:** Formal analysis. **Zhenxing Geng:** Formal analysis. **Yue Qin:** Formal analysis. **Shisong Ma:** Conceptualization, Methodology, Formal analysis, Supervision, Writing – Original draft, Review & Editing, Funding acquisition.

## Declaration of Interests

The authors declare no competing interests.

## Supplemental Information

**Figure S1: Expression of the 23 eye atlas-restricted programs.**

**Figure S2: Expression of selected gene programs in the CRC Visium HD dataset.**

**Table S1: CELLxGENE Census datasets used in the analysis.**

**Table S2: Constituent modules for each gene program.**

**Table S3: The 1,534 gene programs identified for human cells.**

**Table S4: GO enrichment analysis results of the gene programs.**

**Table S5: Annotations of the gene programs.**

**Table S6: Gene programs identified for endothelial cells and other major cell types.**

**Table S7: Gene programs identified for tissues and organs.**

**Table S8: Gene programs identified from the CRC Visium HD sample.**

**Table S9: Annotation of CRC gene programs by comparing to the 1,534 reference programs.**

**Table S10: Enrichment of disease loci within the gene programs.**

## Notes

### Competing Interest Statement

The authors have declared no competing interest.

### Summary of Updates

Added an analysis of batch effects and UMAP visualizations of gene programs (Figure 2; Figure 4B).

## References

Abdulla, S., Aevermann, B., Assis, P., Badajoz, S., Bell, S.M., Bezzi, E., Cakir, B., Chaffer, J., Chambers, S., Cherry, J.M., et al. (2025). CZ CELLxGENE Discover: a single-cell data platform for scalable exploration, analysis and modeling of aggregated data. Nucleic Acids Res 53, D886–d900.

Akwii, R.G., Sajib, M.S., Zahra, F.T., and Mikelis, C.M. (2019). Role of Angiopoietin-2 in Vascular Physiology and Pathophysiology. In Cells.

Andersen, C.L., Christensen, L.L., Thorsen, K., Schepeler, T., Sorensen, F.B., Verspaget, H.W., Simon, R., Kruhoffer, M., Aaltonen, L.A., Laurberg, S., et al. (2009). Dysregulation of the transcription factors SOX4, CBFB and SMARCC1 correlates with outcome of colorectal cancer. Br J Cancer 100, 511–523.

Andrae, J., Gallini, R., and Betsholtz, C. (2008). Role of platelet-derived growth factors in physiology and medicine. Genes Dev 22, 1276–1312.

Barkley, D., Moncada, R., Pour, M., Liberman, D.A., Dryg, I., Werba, G., Wang, W., Baron, M., Rao, A., Xia, B., et al. (2022). Cancer cell states recur across tumor types and form specific interactions with the tumor microenvironment. Nat Genet 54, 1192–1201.

Chaudhary, N., Choudhary, B.S., Shah, S.G., Khapare, N., Dwivedi, N., Gaikwad, A., Joshi, N., Raichanna, J., Basu, S., Gurjar, M., et al. (2021). Lipocalin 2 expression promotes tumor progression and therapy resistance by inhibiting ferroptosis in colorectal cancer. Int J Cancer 149, 1495–1511.

Cheng, S., Li, Z., Gao, R., Xing, B., Gao, Y., Yang, Y., Qin, S., Zhang, L., Ouyang, H., Du, P., et al. (2021). A pan-cancer single-cell transcriptional atlas of tumor infiltrating myeloid cells. Cell 184, 792–809.e723.

D’Andrea, A.D., and Grompe, M. (2003). The Fanconi anaemia BRCA pathway. Nature Reviews Cancer 3, 23–34.

Del Prete, A., Salvi, V., Soriani, A., Laffranchi, M., Sozio, F., Bosisio, D., and Sozzani, S. (2023). Dendritic cell subsets in cancer immunity and tumor antigen sensing. Cell Mol Immunol 20, 432–447.

Elmentaite, R., Kumasaka, N., Roberts, K., Fleming, A., Dann, E., King, H.W., Kleshchevnikov, V., Dabrowska, M., Pritchard, S., Bolt, L., et al. (2021). Cells of the human intestinal tract mapped across space and time. Nature 597, 250–255.

Enright, A.J., Van Dongen, S., and Ouzounis, C.A. (2002). An efficient algorithm for large-scale detection of protein families. Nucleic Acids Res 30, 1575–1584.

Gao, Y., Li, J., Cheng, W., Diao, T., Liu, H., Bo, Y., Liu, C., Zhou, W., Chen, M., Zhang, Y., et al. (2024). Cross-tissue human fibroblast atlas reveals myofibroblast subtypes with distinct roles in immune modulation. Cancer Cell 42, 1764–1783.e1710.

Gavish, A., Tyler, M., Greenwald, A.C., Hoefflin, R., Simkin, D., Tschernichovsky, R., Galili Darnell, N., Somech, E., Barbolin, C., Antman, T., et al. (2023). Hallmarks of transcriptional intratumour heterogeneity across a thousand tumours. Nature 618, 598–606.

Goncharov, N.V., Popova, P.I., Avdonin, P.P., Kudryavtsev, I.V., Serebryakova, M.K., Korf, E.A., and Avdonin, P.V. (2020). Markers of Endothelial Cells in Normal and Pathological Conditions. Biochem (Mosc) Suppl Ser A Membr Cell Biol 14, 167–183.

Goossens-Beumer, I.J., Zeestraten, E.C., Benard, A., Christen, T., Reimers, M.S., Keijzer, R., Sier, C.F., Liefers, G.J., Morreau, H., Putter, H., et al. (2014). Clinical prognostic value of combined analysis of Aldh1, Survivin, and EpCAM expression in colorectal cancer. Br J Cancer 110, 2935–2944.

Hao, Y., Hao, S., Andersen-Nissen, E., Mauck, W.M., 3rd, Zheng, S., Butler, A., Lee, M.J., Wilk, A.J., Darby, C., Zager, M., et al. (2021). Integrated analysis of multimodal single-cell data. Cell 184, 3573–3587.

Hie, B., Bryson, B., and Berger, B. (2019). Efficient integration of heterogeneous single-cell transcriptomes using Scanorama. Nat Biotechnol 37, 685–691.

Hrovatin, K., Sikkema, L., Shitov, V.A., Heimberg, G., Shulman, M., Oliver, A.J., Mueller, M.F., Ibarra, I.L., Wang, H., Ramírez-Suástegui, C., et al. (2024). Considerations for building and using integrated single-cell atlases. Nature Methods.

Ji, L.L., Wei, Y.Z., Jiang, T., and Wang, S.Y. (2014). Correlation of Nrf2, NQO1, MRP1, cmyc and p53 in colorectal cancer and their relationships to clinicopathologic features and survival. Int J Clin Exp Pathol 7, 1124–1131.

Jorstad, N.L., Close, J., Johansen, N., Yanny, A.M., Barkan, E.R., Travaglini, K.J., Bertagnolli, D., Campos, J., Casper, T., Crichton, K., et al. (2023). Transcriptomic cytoarchitecture reveals principles of human neocortex organization. Science 382, eadf6812.

Klötzer, K.A., Abedini, A., Li, S., Balzer, M.S., Liang, X., Levinsohn, J., Ha, E., Dumoulin, B., Hogan, J.J., Quinn, G., et al. (2025). Analysis of individual patient pathway coordination in a cross-species single-cell kidney atlas. Nat Genet 57, 1922–1934.

Kotliar, D., Curtis, M., Agnew, R., Weinand, K., Nathan, A., Baglaenko, Y., Slowikowski, K., Zhao, Y., Sabeti, P.C., Rao, D.A., et al. (2025). Reproducible single-cell annotation of programs underlying T cell subsets, activation states and functions. Nat Methods.

Kotliar, D., Veres, A., Nagy, M.A., Tabrizi, S., Hodis, E., Melton, D.A., and Sabeti, P.C. (2019). Identifying gene expression programs of cell-type identity and cellular activity with single-cell RNA-Seq. eLife 8, e43803.

Kunes, R.Z., Walle, T., Land, M., Nawy, T., and Pe’er, D. (2023). Supervised discovery of interpretable gene programs from single-cell data. Nature Biotechnology.

Kuo, C.F., Grainge, M.J., Zhang, W.Y., and Doherty, M. (2015). Global epidemiology of gout: prevalence, incidence and risk factors. Nature Reviews Rheumatology 11, 649–662.

Lake, B.B., Menon, R., Winfree, S., Hu, Q., Ferreira, R.M., Kalhor, K., Barwinska, D., Otto, E.A., Ferkowicz, M., Diep, D., et al. (2023). An atlas of healthy and injured cell states and niches in the human kidney. Nature 619, 585–594.

Lee, D.D., and Seung, H.S. (1999). Learning the parts of objects by non-negative matrix factorization. Nature 401, 788–791.

Lee, M.Y., Ha, S.E., Park, C., Park, P.J., Fuchs, R., Wei, L., Jorgensen, B.G., Redelman, D., Ward, S.M., Sanders, K.M., et al. (2017). Transcriptome of interstitial cells of Cajal reveals unique and selective gene signatures. PLoS One 12, e0176031.

Li, J., Wang, J., Ibarra, I.L., Cheng, X., Luecken, M.D., Lu, J., Monavarfeshani, A., Yan, W., Zheng, Y., Zuo, Z., et al. (2026). Single-cell atlas of the transcriptome and chromatin accessibility in the human retina. Nature Genetics 58, 418–433.

Lisby, A.N., Flickinger, J.C., Jr., Bashir, B., Weindorfer, M., Shelukar, S., Crutcher, M., Snook, A.E., and Waldman, S.A. (2021). GUCY2C as a biomarker to target precision therapies for patients with colorectal cancer. Expert Rev Precis Med Drug Dev 6, 117–129.

Lopez, R., Regier, J., Cole, M.B., Jordan, M.I., and Yosef, N. (2018). Deep generative modeling for single-cell transcriptomics. Nat Methods 15, 1053–1058.

Montoro, D.T., Haber, A.L., Biton, M., Vinarsky, V., Lin, B., Birket, S.E., Yuan, F., Chen, S., Leung, H.M., Villoria, J., et al. (2018). A revised airway epithelial hierarchy includes CFTR-expressing ionocytes. Nature 560, 319–324.

Morabito, S., Reese, F., Rahimzadeh, N., Miyoshi, E., and Swarup, V. (2023). hdWGCNA identifies co-expression networks in high-dimensional transcriptomics data. Cell Rep Methods 3, 100498.

Ngo, C., Garrec, C., Tomasello, E., and Dalod, M. (2024). The role of plasmacytoid dendritic cells (pDCs) in immunity during viral infections and beyond. Cell Mol Immunol 21, 1008–1035.

Nowicki-Osuch, K., Zhuang, L., Cheung, T.S., Black, E.L., Masqué-Soler, N., Devonshire, G., Redmond, A.M., Freeman, A., di Pietro, M., Pilonis, N., et al. (2023). Single-Cell RNA Sequencing Unifies Developmental Programs of Esophageal and Gastric Intestinal Metaplasia. Cancer Discov 13, 1346–1363.

Oliveira, M.F., Romero, J.P., Chung, M., Williams, S.R., Gottscho, A.D., Gupta, A., Pilipauskas, S.E., Mohabbat, S., Raman, N., Sukovich, D.J., et al. (2025). High-definition spatial transcriptomic profiling of immune cell populations in colorectal cancer. Nat Genet 57, 1512–1523.

Perez, R.K., Gordon, M.G., Subramaniam, M., Kim, M.C., Hartoularos, G.C., Targ, S., Sun, Y., Ogorodnikov, A., Bueno, R., Lu, A., et al. (2022). Single-cell RNA-seq reveals cell type-specific molecular and genetic associations to lupus. Science 376, eabf1970.

Pottel, H., Hoste, L., Dubourg, L., Ebert, N., Schaeffner, E., Eriksen, B.O., Melsom, T., Lamb, E.J., Rule, A.D., Turner, S.T., et al. (2016). An estimated glomerular filtration rate equation for the full age spectrum. Nephrology Dialysis Transplantation 31, 798–806.

Reichart, D., Lindberg, E.L., Maatz, H., Miranda, A.M.A., Viveiros, A., Shvetsov, N., Gärtner, A., Nadelmann, E.R., Lee, M., Kanemaru, K., et al. (2022). Pathogenic variants damage cell composition and single cell transcription in cardiomyopathies. Science 377, eabo1984.

Rigamonti, N., Kadioglu, E., Keklikoglou, I., Wyser Rmili, C., Leow, Ching C., and De Palma, M. (2014). Role of Angiopoietin-2 in Adaptive Tumor Resistance to VEGF Signaling Blockade. Cell Rep 8, 696–706.

Rocha, S.F., Schiller, M., Jing, D., Li, H., Butz, S., Vestweber, D., Biljes, D., Drexler, H.C.A., Nieminen-Kelhä, M., Vajkoczy, P., et al. (2014). Esm1 Modulates Endothelial Tip Cell Behavior and Vascular Permeability by Enhancing VEGF Bioavailability. Circulation Research 115, 581-+.

Ronaldson, P.T., and Davis, T.P. (2024). Blood-brain barrier transporters: a translational consideration for CNS delivery of neurotherapeutics. Expert Opin Drug Deliv 21, 71–89.

Sarrazin, S., Adam, E., Lyon, M., Depontieu, F., Motte, V., Landolfi, C., Lortat-Jacob, H., Bechard, D., Lassalle, P., and Delehedde, M. (2006). Endocan or endothelial cell specific molecule-1 (ESM-1): A potential novel endothelial cell marker and a new target for cancer therapy. Biochimica Et Biophysica Acta-Reviews on Cancer 1765, 25–37.

Schupp, J.C., Adams, T.S., Cosme, C., Raredon, M.S.B., Yuan, Y., Omote, N., Poli, S., Chioccioli, M., Rose, K.-A., Manning, E.P., et al. (2021). Integrated Single-Cell Atlas of Endothelial Cells of the Human Lung. Circulation 144, 286–302.

Shroff, U.N., Gyarmati, G., Izuhara, A., Deepak, S., and Peti-Peterdi, J. (2021). A new view of macula densa cell protein synthesis. Am J Physiol Renal Physiol 321, F689–f704.

Sikkema, L., Ramírez-Suástegui, C., Strobl, D.C., Gillett, T.E., Zappia, L., Madissoon, E., Markov, N.S., Zaragosi, L.-E., Ji, Y., Ansari, M., et al. (2023). An integrated cell atlas of the lung in health and disease. Nature Medicine 29, 1563–1577.

Siletti, K., Hodge, R., Mossi Albiach, A., Lee, K.W., Ding, S.L., Hu, L., Lönnerberg, P., Bakken, T., Casper, T., Clark, M., et al. (2023). Transcriptomic diversity of cell types across the adult human brain. Science 382, eadd7046.

Sollis, E., Mosaku, A., Abid, A., Buniello, A., Cerezo, M., Gil, L., Groza, T., Güneş, O., Hall, P., Hayhurst, J., et al. (2023). The NHGRI-EBI GWAS Catalog: knowledgebase and deposition resource. Nucleic Acids Res 51, D977–d985.

Takeda, A., Salmi, M., and Jalkanen, S. (2023). Lymph node lymphatic endothelial cells as multifaceted gatekeepers in the immune system. Trends Immunol 44, 72–86.

Uribesalgo, I., Hoffmann, D., Zhang, Y., Kavirayani, A., Lazovic, J., Berta, J., Novatchkova, M., Pai, T.P., Wimmer, R.A., László, V., et al. (2019). Apelin inhibition prevents resistance and metastasis associated with anti-angiogenic therapy. EMBO Mol Med 11, e9266.

van der Wijst, M.G.P., Vazquez, S.E., Hartoularos, G.C., Bastard, P., Grant, T., Bueno, R., Lee, D.S., Greenland, J.R., Sun, Y., Perez, R., et al. (2021). Type I interferon autoantibodies are associated with systemic immune alterations in patients with COVID-19. Science translational medicine 13, eabh2624.

Villani, A.-C., Satija, R., Reynolds, G., Sarkizova, S., Shekhar, K., Fletcher, J., Griesbeck, M., Butler, A., Zheng, S., Lazo, S., et al. (2017). Single-cell RNA-seq reveals new types of human blood dendritic cells, monocytes, and progenitors. Science 356, eaah4573.

Wälchli, T., Ghobrial, M., Schwab, M., Takada, S., Zhong, H., Suntharalingham, S., Vetiska, S., Gonzalez, D.R., Wu, R., Rehrauer, H., et al. (2024). Single-cell atlas of the human brain vasculature across development, adulthood and disease. Nature 632, 603–613.

Xu, C., Lopez, R., Mehlman, E., Regier, J., Jordan, M.I., and Yosef, N. (2021). Probabilistic harmonization and annotation of single-cell transcriptomics data with deep generative models. Mol Syst Biol 17, e9620.

Xu, Y., Chen, L., and Ma, S. (2025). SpacGPA: annotating spatial transcriptomes through *de novo* interpretable gene programs. bioRxiv, 2025.2010.2001.679918.

Xu, Y., Wang, Y., and Ma, S. (2024). SingleCellGGM enables gene expression program identification from single-cell transcriptomes and facilitates universal cell label transfer. Cell Rep Methods 4, 100813.

Yan, D., Wiesmann, M., Rohan, M., Chan, V., Jefferson, A.B., Guo, L.D., Sakamoto, D., Caothien, R.H., Fuller, J.H., Reinhard, C., et al. (2001). Elevated expression of *axin2* and *hnkd* mRNA provides evidence that Wnt/β-catenin signaling is activated in human colon tumors. Proceedings of the National Academy of Sciences of the United States of America 98, 14973–14978.

Yang, J., Song, X., Zhang, H., Liu, Q., Wei, R., Guo, L., Yuan, C., Chen, F., Xue, K., Lai, Y., et al. (2024a). Single-cell transcriptomic landscape deciphers olfactory neuroblastoma subtypes and intra-tumoral heterogeneity. Nature Cancer.

Yang, Y., Chen, X., Pan, J., Ning, H., Zhang, Y., Bo, Y., Ren, X., Li, J., Qin, S., Wang, D., et al. (2024b). Pan-cancer single-cell dissection reveals phenotypically distinct B cell subtypes. Cell 187, 4790–4811.e4722.

Zhang, Y., Xiang, G., Jiang, A.Y., Lynch, A., Zeng, Z., Wang, C., Zhang, W., Fan, J., Kang, J., Gu, S.S., et al. (2023). MetaTiME integrates single-cell gene expression to characterize the meta-components of the tumor immune microenvironment. Nat Commun 14, 2634.

Zhao, H., Ming, T.Q., Tang, S., Ren, S., Yang, H., Liu, M.L., Tao, Q., and Xu, H.B. (2022). Wnt signaling in colorectal cancer: pathogenic role and therapeutic target. Mol Cancer 21.

Zheng, L., Qin, S., Si, W., Wang, A., Xing, B., Gao, R., Ren, X., Wang, L., Wu, X., Zhang, J., et al. (2021). Pan-cancer single-cell landscape of tumor-infiltrating T cells. Science 374, abe6474.

