## Supplementary material for "A gene program dictionary of human cells": Figure S1

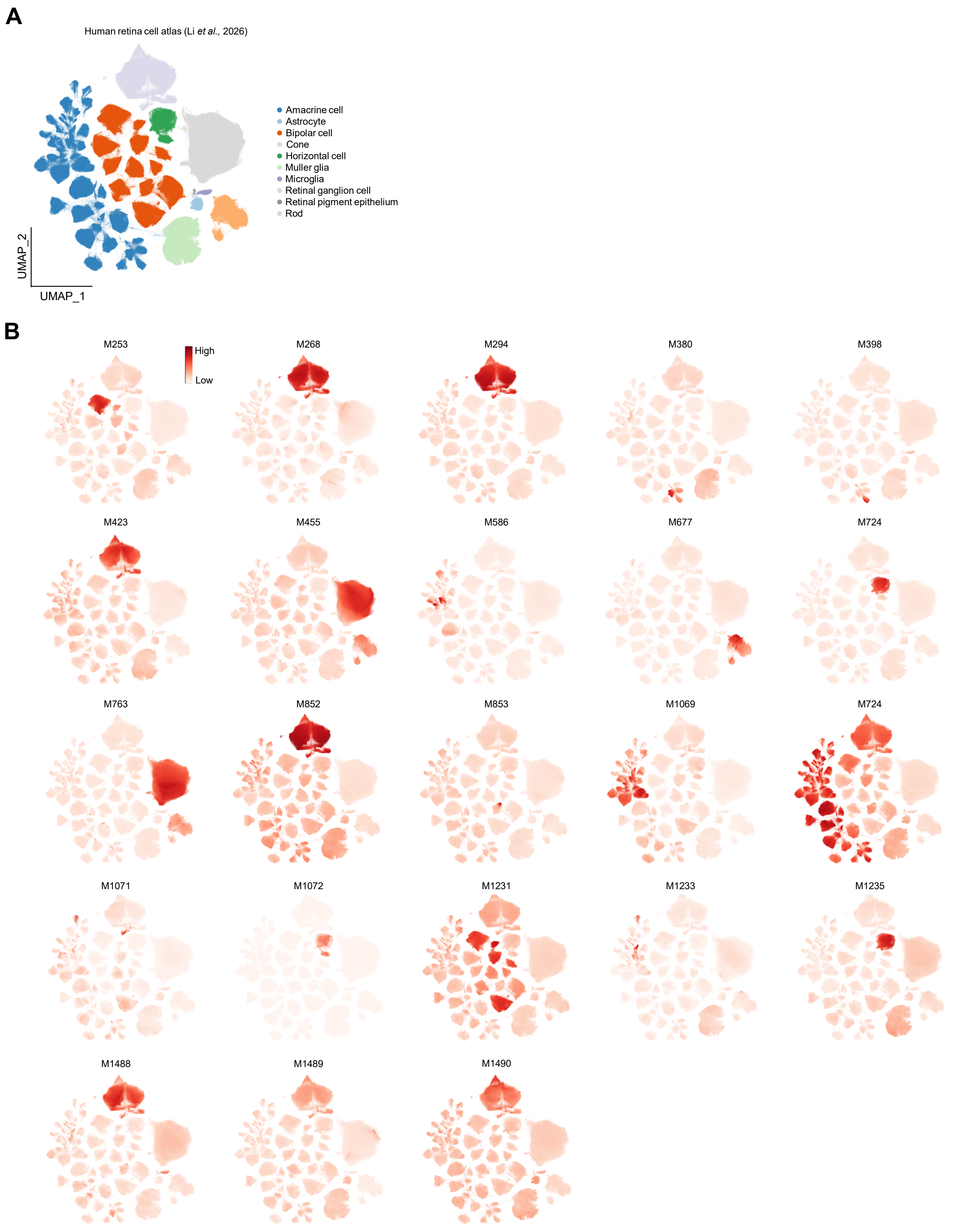


**Figure S1: Expression of the 23 eye atlas-restricted programs.** The programs are visualized based on their expression within the dataset from which they were derived (Li et al. 2026). **(A)** Cell type annotations of the dataset. **(B)** Heatmaps showing the expressio of the 23 programs across cell types.
