## Supplementary material for "A gene program dictionary of human cells": Figure S2

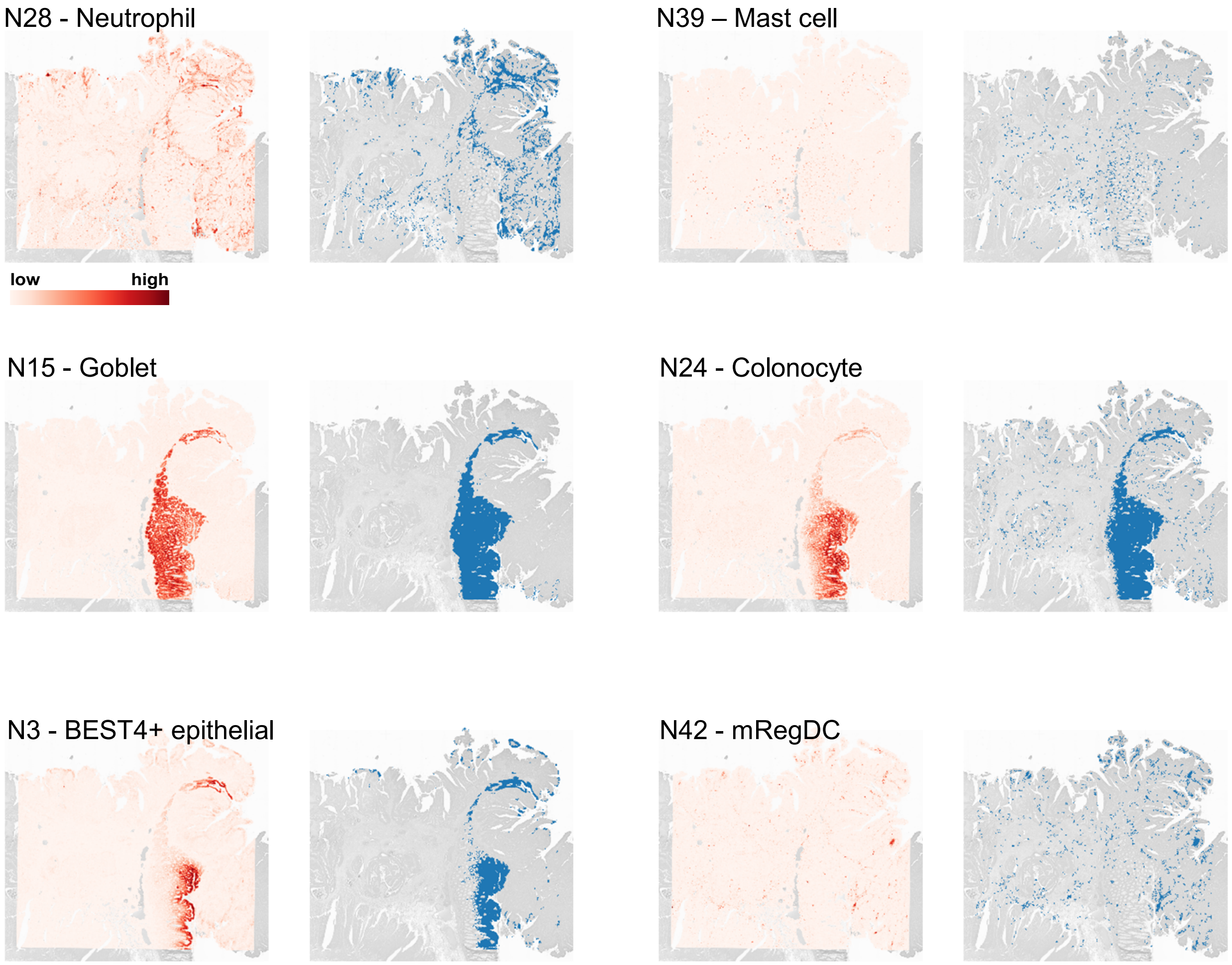


**Figure S2: Expression of selected gene programs in the CRC Visium HD dataset.** For each program, the left panel shows the spatial distribution of its weighted expression across tissue, and the right panel highlights the spots annotated by that program.
